# “Dynamic SUMOylation Controls RNA Polymerase I Extranucleolar Organization and Antigenic Variation in *Trypanosoma brucei*”

**DOI:** 10.64898/2026.08.14.744927

**Authors:** María Agustina Berazategui, Melina Serassio, William Hack, Miguel Navarro, Joana R. Correia Faria, Paula Ana Iribarren, Vanina E. Alvarez

## Abstract

Antigenic variation in *Trypanosoma brucei* relies on strict monoallelic expression of variant surface glycoprotein (VSG) genes from a single telomeric expression site (ES), a process sustained by the extranucleolar RNA polymerase I (Pol I) transcriptional body known as the expression site body (ESB). Although the ESB is essential for VSG expression, the mechanisms governing its assembly and maintenance remain poorly understood. Here, we identify SUMOylation as a central regulator of ESB organization and demonstrate that the balance between SUMO conjugation and deconjugation determines the transcriptional state of VSG expression sites. Ectopic expression of the SUMO protease *Tb*SENP disrupted the highly SUMOylated nuclear focus associated with the active-ES, displaced Pol I from its extranucleolar compartment, and markedly increased VSG *in situ* switching frequency, indicating that continuous SUMOylation is required to preserve ESB integrity. Conversely, targeted recruitment of the SUMO-conjugating enzyme *Tb*UBC9 to a ‘silent’ ES locally restored SUMOylation, induced *de novo* formation of an extranucleolar Pol I compartment, activated transcription of the corresponding telomeric VSG gene, and generated stable antigenic switchers expressing the new surface coat. Local SUMOylation preceded Pol I redistribution, supporting a model in which SUMO-dependent interactions nucleate assembly of a transcriptionally competent ESB. Together, our findings identify SUMOylation as both a structural and regulatory determinant of nuclear organization in *T. brucei* and suggest that dynamic SUMO homeostasis governs the assembly, maintenance, and remodeling of this specialized transcriptional body.

**Significance Statement:** Antigenic variation in *Trypanosoma brucei* depends on the monoallelic expression of Variant Surface Glycoprotein (VSG) genes from a specialized RNA polymerase I transcriptional compartment known as the Expression Site Body (ESB). However, the molecular signals that govern transitions between active and silent expression sites have remained unknown. We show that SUMOylation acts as a reversible molecular switch: disruption of SUMO homeostasis dismantles ESB organization and promotes VSG switching, whereas localized SUMOylation is sufficient to nucleate a functional transcriptional compartment and activate a silent VSG expression site. Our findings establish SUMOylation as a central regulator of nuclear architecture and antigenic variation.

## Introduction

*Trypanosoma brucei* is a protozoan parasite responsible for Human African Trypanosomiasis (sleeping sickness) in humans and Nagana in livestock (1). In the mammalian bloodstream, the parasite resides in the extracellular space, where it must continuously evade the host immune response to establish persistent infections. This ability relies on a sophisticated mechanism of antigenic variation, which allows the parasite to periodically change the identity of its major surface antigen, the Variant Surface Glycoprotein (VSG) (2, 3). The parasite surface is covered by a dense monolayer of VSG molecules that, although highly immunogenic, effectively shield invariant surface proteins from immune detection (4, 5). The *T. brucei* genome encodes over 2,000 *VSG* genes (6, 7). Despite this vast repertoire, expression is strictly monoallelic, with only a single gene expressed at a time from one of approximately twenty subtelomeric expression-sites (*VSG*-ESs) in a strictly monoallelic manner (8). Periodic replacement of the expressed VSG gene - known as VSG switching - generates antigenic diversity within the parasite population, allowing escape from antibody-mediated lysis. VSG switching occurs at low frequency through two major mechanisms: (i) gene conversion or telomere exchange, which replaces the active *VSG* gene by recombination, and (ii) *in situ* switching, in which the active-*VSG*-ES is transcriptionally silenced while a previously ‘silent’ *VSG*-ES becomes activated (9).

In a remarkable deviation from canonical eukaryotic transcription, *VSG* expression is driven by RNA polymerase I (Pol I) (10) from a unique extranucleolar nuclear body termed the Expression Site Body (ESB) (11). The ESB is a dedicated subnuclear compartment that facilitates high-level, processive transcription of the single active-*VSG*-ES, and its formation and maintenance are essential for ensuring monoallelic expression (12). Although the composition of the ESB remained mysterious for over fifteen years following its discovery, recent genetic screens and affinity purifications have begun to unveil its core components. Among the first identified were VSG-Exclusion-Protein-2 (VEX2), an RNA:DNA helicase which regulates monoallelic VSG expression (13), and ESB-Specific-Protein-1 (ESB1), which is involved in transcriptional activation (14). Alongside them is ESBX, which integrates both activities, perhaps by facilitating both ESB1 and VEX2 functions through an unknown mechanism (15). The remaining components - ESB2, ESB3, and ESAP1 - act post-transcriptionally to modulate the polycistronic output of the active VSG ES by downregulating the co-transcribed expression-site-associated genes (ESAGs) to prioritize production of the VSG coat (16). However, the precise molecular mechanisms and functional interplay among these factors remain to be fully elucidated. Notably, the ESB and the chromatin of the active-*VSG*-ES are distinguished by the enrichment of proteins modified by Small Ubiquitin-like modifier (SUMO), suggesting a regulatory role for SUMOylation in *VSG* expression control (17).

SUMOylation is a highly conserved post-translational modification that regulates thousands of proteins involved in gene expression, genome stability, chromosome segregation, and subnuclear organization (18). By dynamically and reversibly modifying functionally related protein networks, the SUMO system coordinates diverse cellular processes, including the assembly of membraneless biomolecular condensates, transcriptional regulation, DNA repair, and cell cycle progression. SUMOylation is mediated by a sequential enzymatic cascade (19). Newly synthesized SUMO precursors are first processed by SUMO-specific proteases (SENPs) to expose the C-terminal Gly-Gly motif essential for conjugation. Mature SUMO is subsequently activated by the E1 enzyme and transferred to the E2-conjugating enzyme UBC9, which can directly modify substrates bearing consensus ΨKxD/E motifs even in the absence of E3 ligases (20). However, SUMO E3 ligases, including the SP-RING, TRIM, and non-canonical families, significantly enhance the efficiency and substrate specificity of SUMO transfer (21). SUMO modification is dynamically controlled by SENPs and other SUMO isopeptidases, including DeSI and USPL1 family members (22), which remove SUMO from target proteins to enable rapid spatiotemporal regulation of protein function. While SENPs generally display broad substrate specificity and participate in multiple steps of the SUMO cycle, DeSI and USPL1 enzymes appear to act on a more restricted set of SUMOylated substrates.

In *T. brucei*, the single SUMO ortholog (*Tb*SUMO) is essential for viability in both bloodstream (BSF) and procyclic (PCF) forms (23, 24), with depletion causing cell cycle arrest, mitotic defects, and abnormal cytokinesis. Proteomic analyses have identified SUMO targets involved in DNA replication and repair, transcription, RNA metabolism, and chromatin remodeling (25). At the active-*VSG*-ES, the SUMO E3 ligase *Tb*SIZ1/PIAS1 establishes a distinctive SUMOylation landscape characterized by high levels of SUMOylated chromatin-associated proteins upstream of the promoter (17). Mechanistically, the largest RNA Pol I subunit (*Tb*RPA1) undergoes SUMOylation and this SUMOylated fraction localizes to an extranucleolar site, as revealed by proximity ligation assays. Functional studies demonstrate that depletion of SUMO-conjugated proteins through *Tb*SUMO, *Tb*UBC9 or *Tb*SIZ1 knockdown significantly reduces RNA Pol I occupancy at the *VSG*-ES, resulting in decreased *VSG* transcript levels. Collectively, these observations support a model in which SUMOylation functions as a positive epigenetic and structural mark that promotes high-level transcription of the active *VSG* gene.

The core SUMO conjugation machinery - comprising *Tb*SUMO, the E1 activating enzyme, and the E2 conjugating enzyme *Tb*UBC9 - has been functionally reconstituted *in vitro* (26) and in heterologous systems (27), validating its biochemical activity. In contrast, considerably less is known about SUMO deconjugation. *Tb*SENP has been shown to process SUMO precursor, deconjugate SUMO-modified substrates, and edit SUMO chains *in vitro* (27), and its depletion leads to increased cellular SUMOylation in BSF parasites (28), consistent with a role in SUMO homeostasis. However, its specific contribution to SUMO-regulated processes and antigenic variation *in vivo* remains unexplored.

In this work, we investigate the consequences of perturbing SUMO dynamics on *VSG* expression. We show that inducible overexpression of the SUMO protease *Tb*SENP disrupts the nuclear SUMOylated focus and promotes *VSG* switching, whereas targeted recruitment of the SUMO conjugation machinery to a ‘silent’ *VSG*-ES activates its transcription. Localized SUMOylation at the promoter of a ‘silent’ *VSG*-ES is sufficient to initiate and sustain transcription along the entire transcription unit, and trigger switching toward the corresponding VSG variant. Together, these findings reveal SUMOylation as a dynamic and reversible post-translational modification that governs antigenic variation in *T. brucei* by coordinating (or integrating) nuclear organization and transcriptional control.

## Results

### Inducible *Tb*SENP ectopic expression promotes global deSUMOylation and impairs parasite proliferation

Despite the established role of SUMOylation in *VSG* transcription, the cellular activity and biological function of the SUMO protease *Tb*SENP (Tb927.9.2220) remains poorly understood. To investigate its role, we generated a BSF cell line with a tetracycline-inducible expression of *Tb*SENP. Multiple independent transfectants were obtained (Supplementary Figure 1), and a clone exhibiting consistent inducible expression with no detectable basal expression was selected for subsequent analyses. Western-blot analysis confirmed inducible *Tb*SENP expression, revealing a protein band of the expected molecular weight (∼85 kDa) together with a lower-molecular-weight form (Figure 1A). Increased *Tb*SENP expression following doxycycline addition was further confirmed by qPCR (Figure 1B). To determine which *Tb*SENP species was associated with deSUMOylating activity *in vivo*, we performed washout experiments following induction. Overexpression resulted in a reduction of high-molecular-weight SUMO conjugates, demonstrating efficient deSUMOylation (Figure 1C and Supplementary Figure 1B and C). Washout experiments revealed that restoration of SUMO conjugates correlated with the disappearance of the ∼85-kDa species, whereas the lower-molecular-weight form remained detectable (Supplementary Figure 1C). These observations suggest that the full-length *Tb*SENP is the catalytically active enzyme *in vivo*, while the smaller species likely represents a processed form with reduced or no detectable deSUMOylating activity.

**Figure 1.**
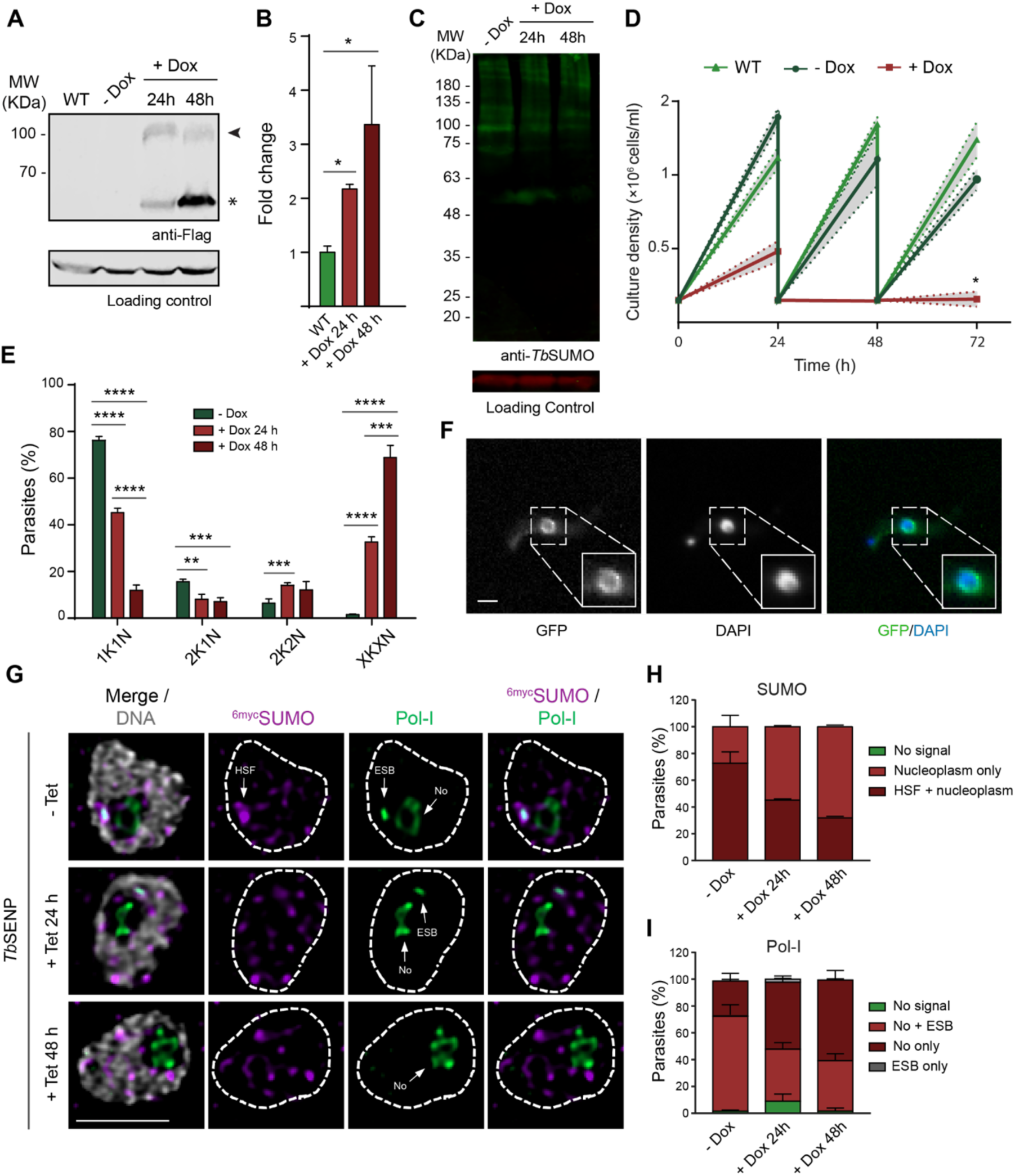
*Tb*SENP overexpression disrupts both the High SUMOylated Focus (HSF) and the Expression-Site-Body (ESB) in *T. brucei* bloodstream forms. (A) Doxycycline inducible expression of *Tb*SENP in BSF parasites leads to a reduction of high-molecular-weight SUMO conjugates. Whole-cell extracts were analysed by Western blot using anti-Flag antibodies. The arrowhead indicates the full-length protein, while the asterisk indicates a lower-molecular-weight form. Tubuline was used as loading control. **(B)** Quantification of RNA transcript levels of *Tb*SENP after 24 h or 48 h induction with doxycycline. Transcript levels were determined by qRT-PCR and normalized against 7SL. Error bars show standard deviation. Statistical analysis: Student’s t-test, *P<0.05. **(C)** Deconjugation activity of *Tb*SENP on SUMO substrates. Whole-cell extracts of wild type (WT), induced (Dox 24 h, Dox 48 h) or uninduced (Dox 0 h) BSF parasites were analysed by Western blot using anti-*Tb*SUMO antibodies. PABP-C was used as loading control. **(D)** Growth curves of wild type (WT), *Tb*SENP induced (+Dox) and *Tb*SENP uninduced (-Dox) parasites. Error areas show standard deviation. Statistical analysis: multiple Student’s t-test, *P<0.05. **(E)** Cell cycle analysis of *Tb*SENP BSF parasites with (+Dox 24 h, +Dox 48 h) or without (-Dox) induction. Cells were stained with DAPI and the number of nuclei (N) and kinetoplasts (K) per cell was quantified. G1 cells (1 kinetoplast and 1 nucleus, 1K1N), G2 cells (2K1N), postmitotic cells (2K2N) and aberrant cells (XKXN). Error bars show standard deviation. Statistical analysis: multiple Student’s t-test, *P<0.05, **P<0.01, ***P<0.001, ****P<0.0001. **(F)** Cellular localization of GFP-*Tb*SENP. Immunofluorescence analysis of BSF parasites after *Tb*SENP induction for 48 hours using anti-GFP antibodies (green). Nuclear and kinetoplast DNA were visualized by DAPI staining (blue). Representative images with magnified views of nuclear regions are shown. Scale bar: 2 µm. **(G-I)** Fluorescence microscopy analysis of Pol-I (**G/H**) and ^6myc^SUMO (**G/I**) localization following *Tb*SENP^3FLAG^ tetracycline (tet)-inducible overexpression. Cells were stained with the following antibodies: anti-myc and anti-*Tb*Pol-I RPA1. DNA was stained with DAPI (grey). The images in **G** were acquired using a Zeiss LSM980 Airyscan 2 and correspond to 3D projections by brightest intensity of 0.1-μm stacks. Scale bar, 2 μm. ESB, expression-site body; HSF, high SUMOylated focus; No, nucleolus. The graphs in **H** and **I** depict mean values of two biological replicates; >100 G1 cells per condition were analysed 24 and 48 h post-induction in comparison with uninduced samples.

While *Tb*SENP induction was initially well tolerated, prolonged expression impaired parasite growth and caused cytokinesis defects, as evidenced by the accumulation of aberrant cells in DAPI-stained populations (Figure 1D and E). These observations are consistent with the essential role of SUMOylation in parasite physiology. Immunofluorescence microscopy revealed that *Tb*SENP localized predominantly at the nuclear periphery (Figure 1F).

### *Tb*SENP promotes disassembly of HSF and the ESB

BSF wild-type cells have an extra-nucleolar Pol I enriched sub-nuclear compartment, the ESB, which harbours the single active-*VSG*-ES and associates with the HSF (17). Since cells depleted of SUMO-conjugated proteins (by either *Tb*SUMO, *Tb*UBC9 or *Tb*SIZ1 knockdown) showed a significant reduction in Pol I occupancy in the *VSG*-ES (analyzed by ChIP using anti-*Tb*RPA1) (17), we examined whether the subnuclear localization of SUMO and Pol I was affected upon *Tb*SENP overexpression. Immunofluorescence analysis revealed a progressive loss of the HSF following *Tb*SENP induction (Figure 1, G-I). The proportion of cells displaying a detectable HSF decreased from 72.6% in non-induced cells to 45.2% and 31.6% after 24 h and 48 h of induction, respectively. A similar trend was observed for the extranucleolar Pol I focus, which was present in 72.5% of non-induced cells but only 41% and 38.2% of cells after 24 h and 48 h of *Tb*SENP induction. Conversely, the proportion of cells exhibiting exclusively nucleolar Pol I staining increased from 26% to 50% and 60.2% over the same period. These results reveal a strong temporal correlation between HSF disassembly and loss of the ESB-associated Pol I focus following global deSUMOylation.

### *Tb*SENP overexpression leads to changes in the VSG coat

The disruption of HSF and ESB integrity following *Tb*SENP induction suggested that altered SUMO dynamics might compromise the mechanisms maintaining monoallelic *VSG* expression. To test this hypothesis, we monitored surface expression of the active VSG221 (also known as VSG-2) variant following transient *Tb*SENP overexpression, using loss of VSG221 (also known as VSG-6) signal as an indicator of potential switching events. Immunofluorescence assays utilizing monoclonal antibodies against VSG221 together with antibodies recognizing alternative VSG variants (i.e. VSG121) enabled detection of coat changes within the population (Figure 2A, +Dox). *Tb*SENP expression was induced for 24 h, after which the inducer was removed and parasites were maintained for an additional 72 h to allow turnover of the pre-existing VSG coat. Importantly, no parasites lacking VSG221 surface expression were detected in the uninduced control (Figure 2A,-Dox). In contrast, transient *Tb*SENP overexpression resulted in the emergence of approximately 31% of cells showing reduced or absent VSG221 surface signal. Among these, ∼9% exhibited complete loss of VSG221 staining with concomitant acquisition of alternative 121 positive VSG signal (Figure 2B). These results demonstrate that transient disruption of SUMO homeostasis is sufficient to promote VSG switching.

**Figure 2.**
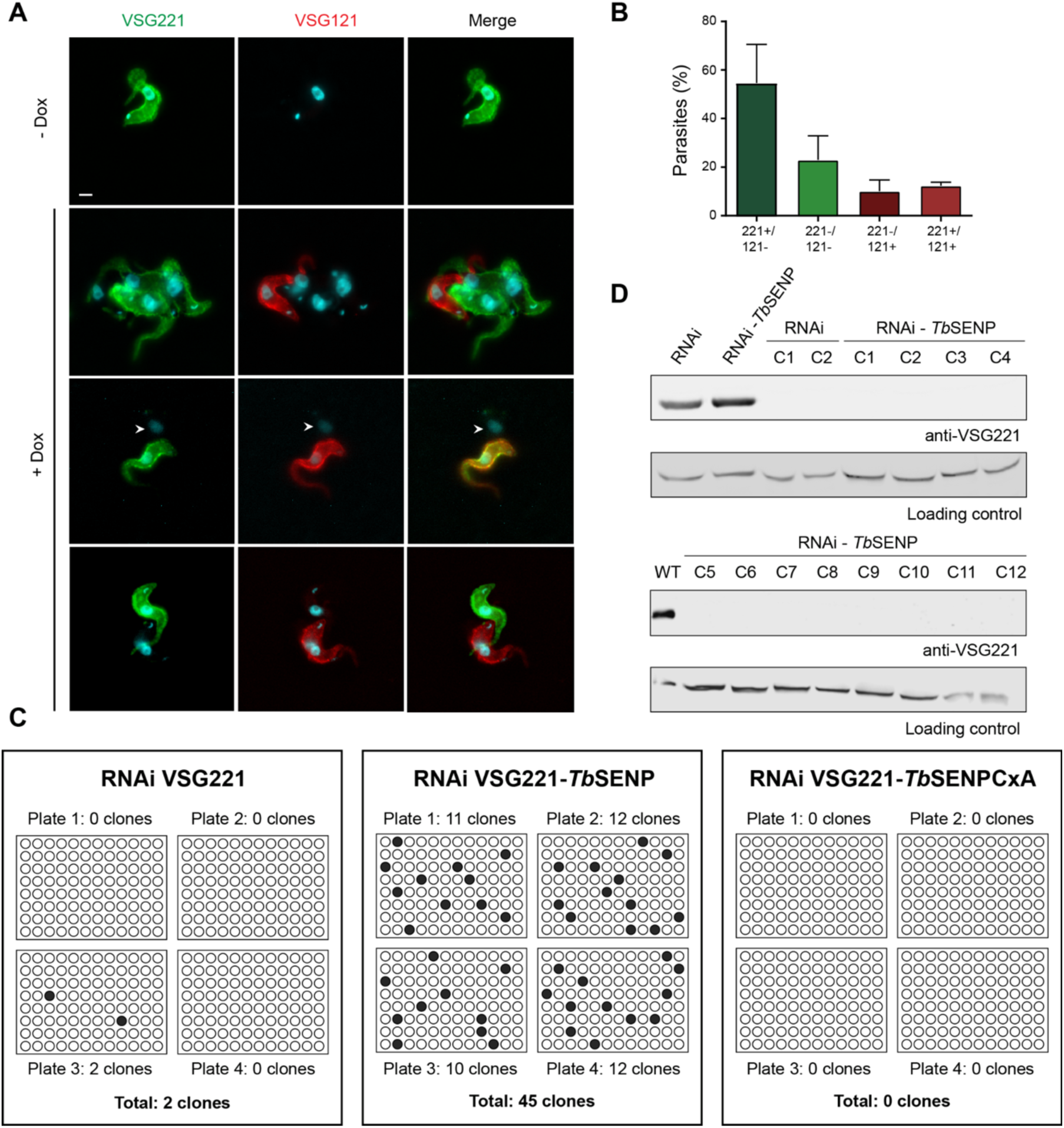
*Tb*SENP transient overexpression leads to increased VSG switching. **(A)** Immunofluorescence analysis of the VSG surface coat in uninduced culture (-Dox) and after 24 h of *Tb*SENP overexpression induction (+Dox), followed by 72 h of growth without inducer to allow VSG coat turnover. The VSG coat was visualized using mouse anti-VSG221 (green) and rabbit anti-VSG121 (red) antibodies. Nuclei and kinetoplasts were stained with DAPI (cyan). White arrowheads indicate parasites potentially expressing a VSG variant distinct from VSG221 or VSG121 (n = 74). Scale bar: 2 µm. **(B)** Quantification of parasite populations according to surface VSG expression. Cells were classified as VSG221+/VSG121-, VSG221+/VSG121+, VSG221-/VSG121+, or VSG221-/VSG121-based on immunofluorescence staining. **(C)** Schematic representation of the plating assay for clones obtained following doxycycline-induced VSG221 RNAi, in the presence or absence of *Tb*SENP or its catalytic mutant (*Tb*SENP CxA). Parasites were seeded in quadruplicate (30 cells/well) with doxycycline. Positive wells showing parasite growth were scored after 7 days. **(D)** Characterization of VSG221 RNAi-resistant parasites obtained after *Tb*SENP overexpression. Whole-protein extracts from 20 randomly selected clones of the RNAi VSG221 (RNAi C1, C2) and RNAi VSG221-*Tb*SENP (RNAi – *Tb*SENP C3-C12) cell line, isolated as described in (A), were analyzed by Western blot using anti-VSG221 antibodies. Wild-type parasites (WT) and uninduced parental RNAi VSG221 (RNAi) and RNAi VSG221-*Tb*SENP (RNAi-*Tb*SENP) cell lines were used as controls. Anti-PABP-C antibodies were used as loading control. Representative clones are shown.

### *Tb*SENP overexpression promotes switching to other telomeric *VSG* variants

To determine whether *Tb*SENP overexpression promotes productive VSG switching, we employed a VSG221 RNAi-based selection system (29) in which only parasites that have switched to an alternative VSG can survive. Simultaneous induction of VSG221 RNAi with either wild-type *Tb*SENP or the catalytically inactive mutant *Tb*SENP(CxA) revealed a striking dependence on enzyme activity: whereas the parental RNAi line yielded only 2 surviving clones, *Tb*SENP overexpression resulted in 45 clones, and no clones were recovered with the inactive mutant (Figure 2C). Western blot and immunofluorescence analysis of randomly selected clones confirmed that all isolated survivors had undergone VSG switching and that survival was not due to RNAi loss (Figure 2D). Sequencing analysis demonstrated that all switch events involved telomeric VSG variants located within expression sites, with VSG800 (MITat 1.18, ES5, 79%) being the most frequently selected, followed by VSG VO2 (MITat 1.9, 14%) and VSG T3 (MITat 1.21, 7%).

### *Tb*SENP overexpression increases switching rates

Since transient disruption of SUMO homeostasis promoted the appearance of alternative VSG variants, we next quantified antigenic switching using an adapted Luria–Delbrück fluctuation assay. Briefly, 20 independent 1 ml cultures of each cell line were initiated at concentrations ranging from 5 to 50 cells ml^−1^ (N_0_). Cells were amplified for 8 generations to reach a final density of 1500 to 15000 cells ml^−1^ (N_t_) and then *Tb*SENP and *VSG* RNAi were induced by adding doxycycline. Each culture was spread over 10 wells of a 96-well plate and wells were allowed to grow for 6–8 days before scoring. The total number of wells were recorded as either switchers or non-switchers, and the switching rates were calculated using the P_0_ method. The estimated switching rates for the RNAi-VSG strain were 1,28×10^-4^ and 4,8×10^-5^, values within the range reported for spontaneous antigenic switching in BSF (∼10⁻⁵ per cell division) (30, 31). In contrast,the *Tb*SENP-overexpressing cell line exhibited a switching rate of 1,36×10^-3^, representing at least a one-order-of-magnitude increase over its parental cell line (Supplementary Table 1).

### Targeted recruitment of *Tb*UBC9 to a silent expression site induces SUMO and Pol I nuclear reorganization

Since global deSUMOylation by *Tb*SENP overexpression increased VSG switching frequency, we asked whether localized SUMO conjugation at a specific *VSG*-ES would be sufficient to trigger its activation. To test this, we used the Lac operator/repressor (LacO-LacI) targeting system developed by Budzak *et al*. (32) to generate a parasite cell line-derived from the HNI VO2 strain (33)-that express an inducible 6xHA-LacI-*Tb*UBC9 fusion protein and carries an array of 50 lacO repeats integrated upstream of the ‘silent’ *VSG221* ES promoter (Figure 3A). Clonal cell lines showed comparable growth rates and expression profiles (Supplementary Figure 2). Induction of 6xHA-LacI-*Tb*UBC9 did not affect parasite growth (Figure 3B) or cell cycle distribution (Figure 3C). Western blot analysis confirmed expression of the fusion protein as early as 8 h post-induction, whereas increased levels of SUMOylated proteins became evident by 48 h post-induction (Figure 3D). Consistent with its targeted recruitment, immunofluorescence analysis revealed its localization in discrete nuclear foci in ∼ 89% of cells (Figure 3E), whereas in cells lacking the LacO repeats it displayed a diffuse nuclear distribution (Supplementary Figure 3C).

**Figure 3.**
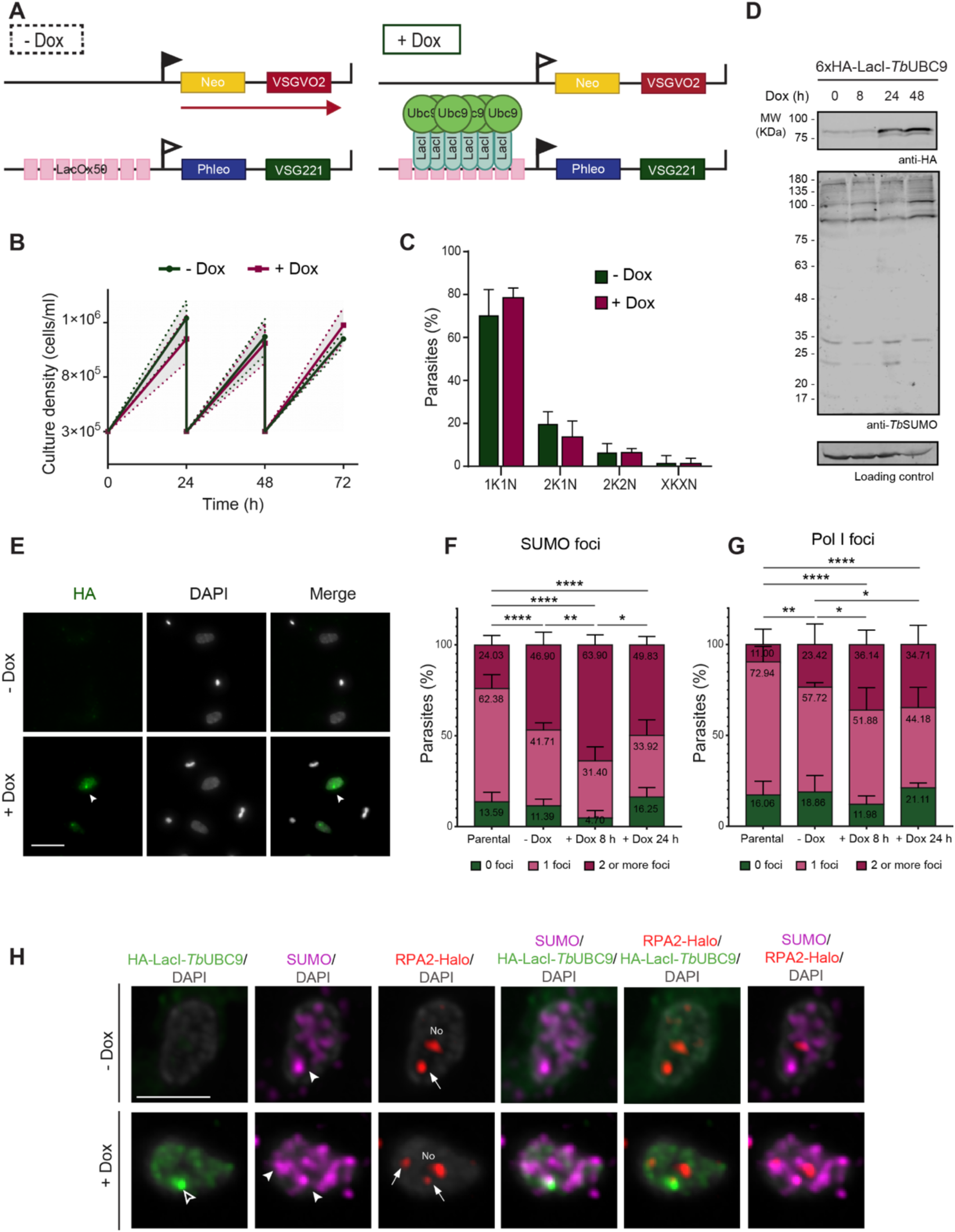
Targeted recruitment of *Tb*UBC9 to a silent expression site drives SUMO and Pol I nuclear reorganization. **(A)** Schematic representation of the recruitment strategy. **(B)** Growth curves of 6xHA-LacI-*Tb*UBC9 cultures with (+Dox) or without (-Dox) doxycycline maintained with 24 hourly subculture (n=3 independent inductions). Error bars show standard deviation (SD). Statistical analysis: multiple Student’s t-test. **(C)** Cell cycle distribution of 6xHA-LacI-*Tb*UBC9 parasites with (+Dox) or without (-Dox) doxycycline induction for 24 h. Cell populations were classified as 1K1N (one kinetoplast, one nucleus), 2K1N (two kinetoplasts, one nucleus), 2K2N (two kinetoplasts, two nuclei), and other (XKXN). Data represent n=3 biological replicates (∼80 parasites per replicate). Error bars show SD. Statistical analysis: multiple Student’s t-test. **(D)** Western blot analysis of 6xHA-LacI-*Tb*UBC9 expression in non-induced (0 h) and doxycycline-induced (+Dox) parasites after 8, 24, and 48 h. Whole cell lysates were probed with anti-HA mAb and anti-*Tb*SUMO antibodies. PABP-C was used as loading control. **(E)** Subcellular localization of 6xHA-LacI-*Tb*UBC9 after 24 h post-induction with (+ Dox) or without (-Dox) doxycycline. Representative images of 1K1N cells immunostained with anti-HA antibody (6xHA-LacI-*Tb*UBC9, green), DAPI (DNA, grey) and merge images are shown. Scale bar: 5 µm. **(F)** Quantification of the percentage of 6xHA-LacI-*Tb*UBC9 cells displaying SUMO foci and **(G)** extranucleolar Pol I foci, after 8 h and 24 h of induction with doxycycline. Data represent three independent experiments (n ≈ 40–60 parasites per replicate). Error bars show SD. Statistical analysis, Chi-square test * P<0.05, **P<0.01, ****P<0.0001 with post-hoc Z-test for the population with two or more foci. Quantification of SUMO foci was performed using an intensity-based threshold (13 AU). **(H)** Subcellular distribution of SUMO and Pol I foci in 6xHA-LacI-*Tb*UBC9 parasites with (+Dox) or without (-Dox) doxycycline induction. Representative images of 1K1N (G1) cells immunostained with anti-HA antibodies (6xHA-LacI-*Tb*UBC9, green), anti-*Tb*SUMO antibodies (HSF, magenta), and JFX554 (Pol I, RPA2-HaloTag, red). Images show merged and individual channels with magnified views of nuclear regions. White arrowheads indicate SUMO foci, arrows Pol I foci and empty arrowhead 6xHA-LacI-*Tb*UBC9. No: nucleolus. Scale bar: 2 µm.

We then asked whether the artificial tethering of the SUMO conjugating enzyme was sufficient to establish *de novo* SUMOylation foci as well as to drive changes in the subnuclear organization of Pol I. Quantitative immunofluorescence analysis revealed a marked increase in the proportion of cells displaying more than two nuclear foci (>2F) for both SUMO conjugates and Pol I following induction of the 6xHA-LacI-*Tb*UBC9 fusion protein (Figure 3F and 3G). For SUMO, the fraction of >2F cells increased from 24.03% in the parental cell line to 46.90% under uninduced conditions (-Dox) consistent with basal expression of the fusion protein. Upon induction, this proportion reached 63.89% at 8 h and subsequently declined to 49.83% at 24 h, indicating a transient accumulation of SUMO foci with maximal nucleation occurring at 8 h. In contrast, Pol I redistribution exhibited a different temporal profile. The proportion of cells containing: >2F Pol I increased progressively from 11.0% in parental cells to 23.42% in the uninduced population, reaching 36.14% and 34.71% after 8 h and 24 h of induction respectively, with the response stabilizing rapidly as no significant change occurred between 8 h and 24 h. Notably, extranucleolar Pol I foci consistently colocalized with SUMO-positive foci in all conditions analyzed, suggesting that Pol I redistribution occurs exclusively at sites enriched in SUMO conjugates (Figure 3H). Together, these results demonstrate that targeted SUMOylation activity at the ‘silent’ *VSG*-ES efficiently nucleates SUMO foci and induces stable redistribution of the Pol I transcription machinery. The distinct temporal profiles of these responses, with SUMO foci peaking before Pol I redistribution stabilizes, suggest that SUMOylation is an early event in this reorganization process.

### Recruitment of *Tb*UBC9 activates transcription of a ‘silent’ *VSG*-ES and promotes VSG switching

Recruitment of 6xHA-LacI-*Tb*UBC9 to the ‘silent’ *VSG221* ES induced transcriptional activation, as assessed by RT-qPCR (Figure 4A). Whereas VSGVO2 transcript levels remained comparable to those of the parental line under uninduced conditions, VSG221 mRNA was already elevated approximately 700-fold, consistent with low-level expression of the fusion protein. Upon induction, VSG221 transcript levels increased further, reaching ∼2000-fold above parental levels at 24 h, concomitant with a marked reduction in VSGVO2 mRNA. By 48 h, VSG221 transcript abundance declined to ∼400-fold, while VSGVO2 expression recovered to near-parental levels. These results indicate that recruitment of *Tb*UBC9 triggers a rapid and robust activation of the ‘silent’ *VSG221*-ES, characterized by an early transcriptional burst followed by a lower but sustained level of expression.

**Figure 4.**
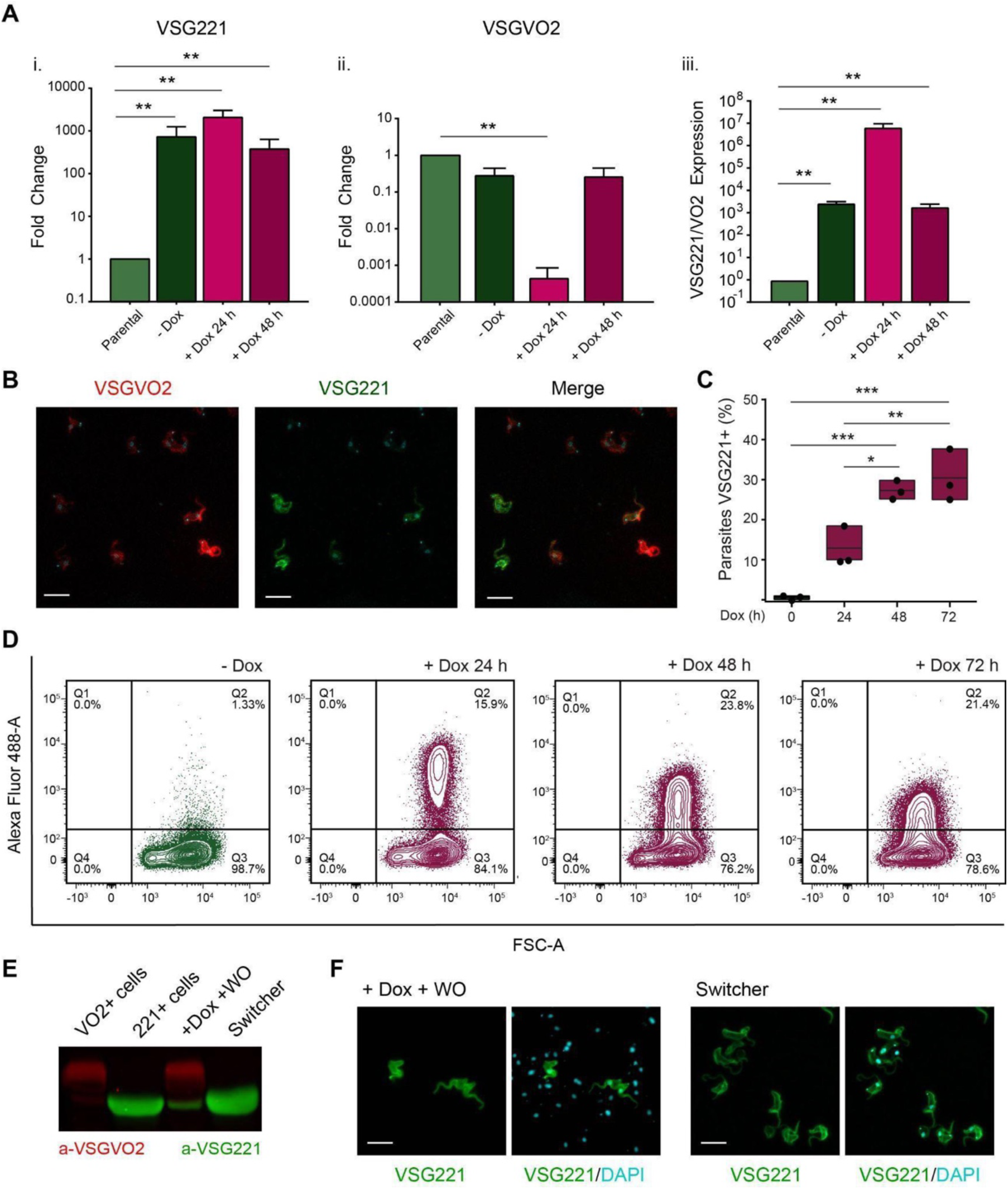
Targeted recruitment of *Tb*UBC9 activates a silent VSG expression site and promotes VSG switching. **(A)** Relative expression analysis of VSG221 and VSGVO2 transcripts in 6xHA-LacI-*Tb*UBC9 parasites. Fold change of VSG221 (i) and VSGVO2 (ii) transcript levels relative to the parental cell line control (HNIVO2+) showing the mean of n = 3 biological replicates under non-induced conditions (−Dox) and after 24 h and 48 h of induction. (iii) Relative mRNA levels of VSG221 with respect to VSGVO2 levels. C1 transcript was used as housekeeping. Error bars show SD. Statistical analysis was performed using a one-sample t-test against the parental cell line, **P<0.01. **(B)** Indirect immunofluorescence of 6xHA-LacI-*Tb*UBC9 parasites using anti-VSG221 mAb (green) and anti-VSGVO2 polyclonal serum (red). A representative image of the population following induction of 6xHA-LacI-*Tb*UBC9 is shown. Nuclei and kinetoplasts were stained with DAPI (cyan). Scale bar: 10 µm. **(C)** Quantification of VSG221+ parasites of 6xHA-LacI-*Tb*UBC9 cell line by indirect immunofluorescence using anti-VSG221 antibodies. Data from n = 3 independent inductions (∼300 parasites per sample) are shown for each time point (Dox 24 h, 48 h, 72 h). The non-induced cell line was used as control (Dox 0 h). Error bars show SD. Statistical analysis was performed using one-way ANOVA, *P<0.05, **P<0.01, ***P<0.001. **(D)** Quantification of switching frequency of 6XHA-LacI-*Tb*UBC9 parasites by flow cytometry. VSG221+ parasites were detected using anti-VSG221 antibodies and anti-mouse AF488 secondary antibodies. A representative example from n = 3 independent inductions is shown (∼100000 events per sample). **(E)** Analysis of VSG221 and VSGVO2 expression by Western blot. VO2+ cells (parental HNIVO2+ cell line) and 221+ cells (HNI221+ cell line) were used as control. +Dox +WO: 6XHA-LacI-*Tb*UBC9 cells induced for 72 h with doxycycline and subsequently maintained in culture for ∼20 days in the absence of inducer; Switcher: clone isolated after ∼20 days washout. Detection was performed using anti-VSG221 (green) and anti-VSGVO2 antibodies (red). **(F)** Indirect immunofluorescence of +Dox +WO and the isolated switcher clone shown in (E), using anti-VSG221 antibodies (green). Nuclei and kinetoplasts were stained with DAPI (cyan). Scale bar: 10 µm.

We next asked whether this transcriptional activation was sufficient to promote productive VSG switching. Indirect immunofluorescence and flow cytometry demonstrated a progressive increase in the proportion of VSG221-positive parasites, from 1.33% in the uninduced cell line to 15.9% at 24 h and 23.8% at 48 h, before stabilizing at 21.4% by 72 h (Figure 4B, C and D). Notably, changes in VSG221 surface expression mirrored the transcriptional profile of the activated *VSG*-ES. The compact VSG221-positive population observed at 24 h corresponded to the peak of VSG221 mRNA accumulation, whereas the broader distributions detected at 48 h and 72 h occurred after transcript levels had declined, likely due to the long half-life of surface VSG molecules.

Importantly, VSG switching depended on the recruitment of 6xHA-LacI–*Tb*UBC9 to the targeted *VSG*-ES, as induction of the fusion protein in the parental cell line lacking lacO repeats upstream of BES1 did not result in comparable switching frequencies (Supplementary Figure 3). Although VO2-negative parasites were detected at frequencies of 3–5% at all time points, VSG221 switchers were only rarely observed in this control. Collectively, these results demonstrate that localized recruitment of the SUMO-conjugating enzyme, rather than its generalized overexpression, is sufficient to promote transcriptional activation of a targeted ‘silent’ *VSG*-ES, leading to productive VSG switching. Together with the preceding observations, these findings support a model in which localized SUMOylation functions as an early event in the establishment of a transcriptionally active-*VSG*-ES.

Finally, to determine whether these events reflected stable antigenic switching rather than transient expression of the new VSG, VSG221-positive parasites were isolated from cultures in which 6xHA-LacI-*Tb*UBC9 expression had been induced for 72 h and propagated for approximately 20 days in the absence of doxycycline. Western blot and immunofluorescence analyses demonstrated uniform VSG221 expression throughout the isolated clones, indicating that recruitment of 6xHA-LacI-*Tb*UBC9 induces productive and heritable VSG switching that is maintained following withdrawal of the inducing stimulus.(Figure 4E and F).

## Discussion

Antigenic variation in *T. brucei* depends on the strict monoallelic expression of *VSG* genes from a single telomeric ES. This process is mediated by a specialized extranucleolar transcriptional body, the ESB, which concentrates Pol I and associated factors to sustain high-level *VSG* transcription (11). However, the molecular mechanisms governing the establishment and maintenance of this unique nuclear compartment remain largely unknown. Here, we identify SUMOylation as a key determinant of Pol I subnuclear organization and show that the dynamic balance between SUMO conjugation and deconjugation regulates the formation and stability of this specialized transcriptional body.

Transcription of the active *VSG* gene is orchestrated by a set of specialized activators that operate within a highly SUMOylated nuclear environment (17). The SUMO E3 ligase *Tb*SIZ1 promotes SUMO conjugation at the active-*VSG*-ES, where several positive regulators of *VSG* transcription either undergo SUMO modification or are predicted to engage in SUMO-dependent interactions. For example, the chromatin remodeler SNF2PH requires SUMOylation at specific N-terminal residues for its recruitment to the active *VSG* promoter and the ESB (34). Likewise, ESB1, a bloodstream stage-specific factor essential for high-level transcription and Pol I recruitment (14), contains predicted SUMO-interacting motifs (SIMs) together with an N-terminal RING/U-box-like domain, a structural feature shared by many ubiquitin and SUMO E3 ligases. Although ligase activity has not been demonstrated for ESB1, these features raise the intriguing possibility that it participates in SUMO-dependent organization of the ESB. Similarly, ESBX, another factor required for *VSG*-ES transcription, and VEX2, an indispensable factor for strict monoallelic *VSG* expression, both required for Pol I extra-nucleolar localization (15, 35), are also predicted to contain SIMs. Moreover, ESB2 and ESAP1, involved in negative regulation of *ESAG* levels at the active-*VSG*-ES, also contained predicted SIMs. Collectively, these observations suggest that the active-*VSG*-ES is enriched in proteins capable of participating in a network of SUMO–SIM interactions. Such multivalent interactions could provide the molecular framework required for the assembly and maintenance of the ESB. In this model, SUMOylation would promote the stable association of transcription factors, chromatin remodelers and Pol I, effectively acting as a “molecular glue” that reinforces the transcriptional machinery. Moreover, localized SUMOylation could initiate a self-reinforcing positive feedback mechanism during the assembly process, whereby recruitment of SUMOylated proteins and SIM-containing partners promotes further SUMO conjugation, progressively amplifying the local interaction network and stabilizing the transcriptional compartment.

SUMOylation is established as a master regulator of the subnuclear landscape across diverse eukaryotic models, orchestrating the spatial organization of the nucleus through the “group modification” of functionally related protein complexes (18). A defining mechanism for this structural role is the assembly of biomolecular condensates, a process extensively characterized in mammalian cells through the formation of PML nuclear bodies (36). Beyond these discrete structures, SUMOylation is essential for high-order chromatin organization and the maintenance of cellular identity; for instance, in mammalian embryonic stem cells, it safeguards pluripotency by enforcing repressive states on the two-cell (2C) embryo-like program through the modification of PRC1.6 and SETDB1 (37). This role in heterochromatin stability is further conserved in fission yeast and mammals, where the modification of HP1 is critical for its initial targeting and subsequent propagation (38). Furthermore, the SUMO system ensures genomic stability by coordinating conserved chromosome dynamics - such as Topoisomerase II modification for centromeric cohesion in yeast and vertebrate models - and by driving the relocation of heterochromatic DNA breaks for specialized repair at the nuclear periphery, as observed in yeast and Drosophila (39). Ultimately, the ability of SUMOylation to act as a molecular scaffold provides the precise spatial compartmentalization required to manage complex nuclear functions across the eukaryotic lineage.

Building upon these established principles, we propose that similar mechanisms govern the assembly of the ESB in *T. brucei*. The exceptionally high local concentration of SUMO conjugates at the active-*VSG*-ES is consistent with this idea and further supports the view that the ESB together with the Highly SUMOylated Focus (HSF) may constitute a phase-separated transcriptional condensate.

Overexpression of the SUMO protease *Tb*SENP disrupted the HSF, caused delocalization of Pol I from its extranucleolar site, and increased VSG switching frequency - likely through *in situ* activation of ‘silent’ *VSG*-ESs. These observations suggest that SUMOylation is required to maintain the nuclear architecture necessary for stable expression site activity. Intriguingly, the phenotype resulting from HSF disruption parallels that observed upon cohesin impairment (12). Just as cohesins ensure the prolonged association of replicated sister chromatids with the ESB until chromosome segregation, tight regulation of SUMO protease activity may be essential for preserving HSF integrity throughout the cell cycle. Such regulation would maintain a stable transcriptional environment during interphase while permitting controlled remodeling at the end of mitosis, thereby contributing to the faithful epigenetic inheritance of the active-ES state in daughter cells.

SUMO homeostasis is maintained through a dynamic balance between SUMO conjugation by the E1-E2-E3 enzymatic cascade and deconjugation by SUMO proteases. Regulation of SUMO protease abundance, localisation and catalytic activity has emerged as an important mechanism for controlling SUMO homeostasis in diverse eukaryotes. Whether SUMO homeostasis in *T. brucei* is similarly regulated through modulation of *Tb*SENP remains unknown. Published transcriptomic datasets indicate that the *Tb*SENP transcript is present at low steady-state abundance but is not unusually unstable, exhibiting a half-life of approximately 23 min in bloodstream forms and 13 min in procyclic forms (40). Likewise, quantitative proteomic analyses detect *Tb*SENP at relatively low abundance (41), although information on its protein turnover is currently unavailable. The extensive deSUMOylation and marked impairment of parasite proliferation following ectopic *Tb*SENP expression indicate that maintaining appropriate *Tb*SENP activity is critical for SUMO homeostasis and parasite viability. Furthermore, ectopic *Tb*SENP expression generated two protein species, with restoration of SUMO conjugates correlating with the disappearance of the full-length protein while the lower molecular weight species remains detectable. These findings raise the possibility that *Tb*SENP is regulated post-translationally. Although the lower-molecular-weight species remains uncharacterised, it may represent a proteolytically processed form of *Tb*SENP. Future studies will be required to determine whether *Tb*SENP activity is regulated through modulation of its abundance, protein turnover, subcellular localisation, post-translational modification or proteolytic processing.

Studies in yeast and mammalian cells have revealed that SUMO proteases are subject to tight spatial and temporal regulation during the cell cycle. The yeast protease Ulp1, as well as mammalian SENP1 and SENP2, are predominantly associated with the nuclear pore complex during interphase but relocalize to mitotic structures during cell division. In particular, SENP1 and SENP2 accumulate at kinetochores, where they regulate key mitotic events, including chromosome alignment and sister chromatid segregation (42). Likewise, Ulp1 is required for proper G2/M progression in yeast (43). Together, these observations highlight that both the abundance and localization of SUMO proteases must be tightly controlled to ensure faithful chromosome segregation and cell cycle progression. Immunofluorescence microscopy revealed that ectopically expressed *Tb*SENP localized predominantly at the nuclear periphery, consistent with its previously reported nuclear pore-associated localisation in the TrypTag database, its enrichment in the NUP110 interactome (44), and recent proximity-labeling studies predicting localisation at the nuclear basket and nuclear side of the nuclear pore complex (45). Interestingly, overexpression of *Tb*SENP in *T. brucei* has also been associated with cytokinesis defects, suggesting that excessive deSUMOylation may perturb cell division in this parasite. In addition, the condensin subunit SMC4 is SUMOylated in *T. brucei* (25), raising the possibility that SUMO-dependent regulation of chromosome architecture and segregation may contribute to the phenotypes observed upon *Tb*SENP overexpression.

Conversely, targeted recruitment of the SUMO-conjugating enzyme *Tb*UBC9 to a ‘silent’ *VSG*-ES was sufficient to locally re-establish SUMOylation and activate transcription of the corresponding *VSG* gene, likely through recruitment of Pol I. Notably, recruitment of *Tb*UBC9 first induced a rapid increase in cells displaying multiple SUMO-enriched nuclear foci, which peaked at 8 h, whereas redistribution of extranucleolar Pol I occurred more gradually and remained stable thereafter. These distinct temporal profiles suggest that localized SUMOylation represents an early event that precedes and facilitates reorganization of the Pol I transcription machinery during activation of a ‘silent’ *VSG*-ES. Consistent with this model, the increase in cells containing multiple Pol I-enriched nuclear foci coincided with the significant rise in *VSG221* transcript levels observed at 24 h, indicating that *de novo* Pol I compartment formation is closely associated with productive activation of the ‘silent’ *VSG*-ES. The transient detection of cells harboring two Pol I-enriched nuclear foci is consistent with previous studies reporting that two transcriptionally active Pol I compartments can coexist during the establishment of a new active expression site (46), supporting the notion that our tethering approach recapitulates key intermediates of the natural switching process.

These findings distinguish localized SUMOylation from previous perturbations of VSG expression-site regulators, including VEX2 depletion (13), ESB1 overexpression (14) and ESBX depletion (15), which have been proposed to generate an enhanced “trickle” transcriptional state without efficient establishment of a fully engaged expression site. One explanation, supported by recent single-cell RNA-seq analyses following VEX2 depletion (35), is that simultaneous activation of multiple VSG expression sites results in competition for limiting transcriptional and RNA-processing factors, preventing any individual expression site from maturing into a fully functional transcriptional state. In contrast, localized SUMOylation promoted the transition to a fully engaged expression-site state, supporting processive transcription of the corresponding telomeric *VSG* gene, replacement of the surface VSG coat and the generation of stable switchers. To our knowledge, this is the first demonstration that targeted recruitment of a regulatory activity to a defined ‘silent’ *VSG*-ES is sufficient to generate a new functional Pol I transcriptional compartment that culminates in stable replacement of the surface VSG coat and establishment of a heritable monoallelic *VSG* expression state.

Interestingly, although *VSG221* transcript levels subsequently declined following prolonged LacI-*Tb*UBC9 induction, VSG221-positive parasites persisted and continued to accumulate, consistent with the long half-life and slow turnover of the VSG surface coat. At the later time points, however, the VSG221-positive population exhibited broader fluorescence distributions and lower mean fluorescence intensities, indicating increased heterogeneity in VSG221 surface abundance. One possible explanation is that localized SUMOylation efficiently initiates activation of a ‘silent’ VSG expression site, but only a subset of parasites successfully completes the transition to a fully stable transcriptional state. Commitment to VSG switching likely requires coordinated recruitment of additional transcriptional and chromatin-associated factors beyond the initial SUMO-dependent activation event. Parasites that fail to complete this transition may only transiently activate *VSG221*, resulting in reduced or heterogeneous surface expression despite initial induction. By contrast, parasites that complete the switch would be expected to establish a single mature ESB associated with the newly active expression site. Alternatively, persistent tethering and prolonged overexpression of *Tb*UBC9 may maintain local SUMOylation beyond its physiological level and temporal window, promoting continued remodelling of the transcriptional machinery or repeated nucleation of SUMO-enriched components, impairing stabilisation of the newly activated expression site in a subset of parasites.

The dynamic interplay between SUMO conjugation and deconjugation emerges as a central mechanism regulating monoallelic *VSG* expression. Excessive deSUMOylation upon *Tb*SENP induction leads to the loss of ESB integrity, suggesting that SUMOylation must be continuously maintained to preserve the active ES. Conversely, enforced SUMO conjugation at a ‘silent’ *VSG*-ES can override epigenetic silencing and trigger transcriptional activation, providing the nucleation event that initiates assembly of a transcriptionally competent Pol I compartment possibly through a self-reinforcing SUMO–SIM interaction network. These findings imply that SUMO homeostasis defines the transcriptional state of each ES and that fluctuations in SUMOylation may underlie natural *in situ* switching events. In this model, transient reductions in SUMO modification could drive ESB disassembly, allowing the establishment of a new SUMO-enriched focus at an alternative ES, thereby promoting the exchange of active and silent VSGs. This dynamic mechanism offers an elegant solution to the challenge of monoallelic control: SUMOylation both establishes and stabilizes the active transcriptional compartment, while deSUMOylation provides a means to reset the system. The SUMO pathway thus integrates structural and regulatory functions, linking post-translational modification with nuclear organization and gene expression plasticity.

## Materials and Methods

### Trypanosome culture

The monomorphic bloodstream form *T. brucei brucei* 427 MITat 1.2 single-marker cell line (47), constitutively expressing an ectopic copy of T7 RNA polymerase and tetracycline repressor, was used in this work. Recruitment experiments were performed using the BSF *T. brucei brucei* 427 HNI MITat 1.9 strain (33). Parasites were cultured in HMI-9 media (Life Technologies, Carlsbad, CA, USA) containing 10% (v/v) heat-inactivated fetal bovine serum (Natocor, Córdoba, Argentina) and incubated at 37 °C with 5% CO_2_ in presence of 2.5 µg/mL G418 or 5 µg/mL Hygromicine and 0.1 µg/mL Puromicine (InvivoGen, San Diego, CA, USA).

Growth curves were performed by triplicate seeding 10^5^ parasites/ml in 10 ml final volume and counted daily using a Neubauer chamber.

Growth curves of 6xHA-LacI-*Tb*UBC9 parasites were performed in triplicate by seeding 0.3×10^6^ parasites/ml and counting daily using a Neubauer chamber, with 24h subculture to the initial density.

### Transgenic cell lines generation

To generate an inducible *Tb*SENP expressing cell line, *Tb*SENP ORF (*Tb*927.9.2220) was amplified by PCR from *T. brucei* genomic DNA using 5’-CCCGGGTATGGCAGATATCCTTTTAAATGCCG-3’ as forward primer and 5’-ACTAGTCGCCTTCGTGTATAAGGCCAGT-3’ as reverse primer. The amplification products were cloned into pGEM-T Easy vector (Promega, Madison, WI, USA), fused to a GFP or tri-Flag N-terminal epitope and sequenced (Macrogen, Seoul, Korea). These constructions were then cloned into a modified pLew100v5 with resistance to blasticidine using *Xba*I and *Nhe*I restriction sites. The vector was linearized with *Not*I before transfection in SM cell line or expressing trypanosomes. TriFlag-*Tb*SENP active site mutant was generated by PCR-based site-directed mutagenesis to replace the catalytic cysteine by an alanine residue and cloned into pLew100v5 as previously described using 5’-CAGAATTCGTCTGATGCGGGTGTTTTTGTGTG-3’ as fwd primer and 5’-CACACAAAAACACCCGCATCAGACGAATTCTG-3’ as rv primer.

For epitope tagging at the native locus (48) a plasmid (pNAT^TAGx^ *BSD*) was generated to add an N-terminal 6× c-myc to SUMO. A 672 bp amplicon containing the SUMO open reading frame and downstream untranslated region was cloned via AvrII/BamHI. The plasmid was linearized with NdeI prior to transfection. TbSUMO_NTagging 5’-AGCTCCTAGGATGGACGAACCCACTCATAA-3’ was used as fwd primer and TbSUMO_NTagging 5’ - AGCTGGATCCGTCCCGTTCCAACCATAAAC-3’ as rv primer.

For generating the VSG221-RNAi cell line, a 803 bp fragment of VSG 221 (position 122 to 925 of GenBank accession X56762), was cloned into a p2T7 vector with resistance to phleomycin using *Bam*HI and *Spe*I restriction sites (49). The vector was linearized with *Not*I and used to transfect SM parasites. RNAi-VSG221-*Tb*SENP cell line was obtained by transfecting TriFlag-*Tb*SENP pLew100v5 into VSG221-RNAi cell line.

The LacO 6xHA-LacI-TbUBC9 cell line was generated from the bloodstream-form *T. brucei* HNI 221+ parental strain, in which the VSG221 BES1 is active and carries the hygromycin resistance cassette, whereas the silent VO2 BES2 contains a neomycin resistance cassette. To generate the HNI-LacO cell line, an array of 50 lac operator (LacO) repeats linked to a phleomycin resistance cassette was integrated immediately upstream of the active VSG221 BES1 promoter by transfection of the pBES1-LacOx50 plasmid linearized with AvrII and KpnI (32). Correct integration was selected with phleomycin and confirmed by PCR. To visualize RPA2 localization, the Halo-RPA2-Hyg construct was transfected into the HNI-LacO cell line, generating the LacO Halo-RPA2 cell line. Stable transformants were selected with hygromycin.

To generate clones in which the VO2 BES2 was active, antigenic switching was induced by culturing LacO Halo-RPA2 parasites for at least four days in the absence of phleomycin, allowing loss of selective pressure on the LacO-containing VSG221 BES1. Parasites were subsequently cloned by limiting dilution in 96-well plates in HMI-9 medium containing G418 to select for activation of the VO2 BES2. Individual clones were screened by Western blot analysis for loss of VSG221 expression and acquisition of VSGVO2 expression. Clones exhibiting the expected VSG expression profile were used as the LacO VO2+ Halo-RPA2 cell line.

Finally, to generate inducible expression of the 6xHA-LacI-*Tb*UBC9, the fusion protein coding sequence was cloned into the p*Tb*FIX expression vector (50), in which transgene expression is driven by a tetracycline-inducible rRNA promoter. The vector additionally contains a constitutively expressed dicistronic transcription unit encoding a nucleus-targeted tetracycline repressor and the puromycin resistance gene, allowing regulation of transgene expression and selection of stable transformants with puromycin. The resulting plasmid was transfected into the LacO VO2+ Halo-RPA2 cell line to generate the LacO 6xHA-LacI-*Tb*UBC9 cell line.

The expression of *Tb*SENP, VSG221 RNAi or 6xHA-LacI-*Tb*UBC9 parasites was induced by addition of 1 μg/ml of doxycycline (Sigma, Saint Louis, MO, USA) to the culture media.

### Transfection

Approximately 2×10^7^ BSF trypanosomes in log phase (∼1.0×10^6^ cells/mL) were collected by centrifugation (1000 g for 10 minutes) at room temperature. The cell pellet was resuspended in 90 μl of transfection buffer (90 mM NaH_2_PO₄/Na_2_HPO₄, 5 mM KCl, 0.15 mM CaCl₂, and 50 mM HEPES pH 7.3). Subsequently, 10 μl of linearized DNA (5–10 μg) was added, and the mixture was transferred to a 2-mm electroporation cuvette (BTX Harvard Apparatus,Holliston, MA, USA). Electroporation was performed using an Amaxa Nucleofector with program X-001. Following electroporation, parasites were resuspended in 10 ml of antibiotic-free HMI-9 medium and incubated under standard growth conditions for 6 h. Cells were then diluted in culture medium containing the appropriate selection antibiotics and distributed into a 24-well plate at 1 ml per well (approximately 2 × 10⁵ parasites ml⁻¹). Plates were incubated at 37°C with 5% CO₂ until resistant parasites were detected.

### Western blot analysis

Cell extracts were resuspended in Laemmli’s sample buffer (0.125 M Tris pH 6.8, 4% (w/v) SDS, 20% (v/v) glycerol, 100 mM DTT) and boiled for 5 min. Samples were resolved by SDS-PAGE and transferred to a nitrocellulose Hybond ECL membrane (GE Healthcare, Pittsburgh, PA, USA) and probed with mouse anti-Flag M2 mAb 1:1000 (Sigma), mouse *Tb*SUMO polyclonal antibody 1:500 (51, 52), rat anti-HA mAb 1:500 (Roche, Basel, Switzerland), mouse anti-VSG221 mAb 1:500, rabbit anti-VSG VO2 polyclonal antibody 1:1000, mouse anti-Actin 1:1000, mouse anti α-tubulin clone B-5-1-2 (Sigma), rabbit anti-PABPc 1:1000 (53) or rabbit anti-BiP 1:5000. Alexa Fluor 790 AffiniPure goat anti-mouse IgG (H+L), Alexa Fluor 680 AffiniPure goat anti-rabbit IgG (H+L), Alexa Fluor 680 AffiniPure goat anti-mouse, Alexa Fluor 790 AffiniPure goat anti-rat and Alexa Fluor 800 AffiniPure goat anti-Rabbit secondary antibodies (Jackson Immunoresearch Laboratories, West Grove, PA, USA), diluted 1:20000 were detected using an Odyssey laser-scanning system (LI-COR Biosciences, Lincoln, NE, USA).

### Immunofluorescence

Parasites were fixed with 1% paraformaldehyde in PBS for 20 min, permeabilized with 0.1% Triton X-100 10 min, followed by incubation in 3% BSA for 30 min for blocking. Mouse anti-*Tb*SUMO polyclonal antibody 1:250, mouse anti-GFP mAb 1:1000 (Sigma), mouse anti-VSG221 mAb 1:250, rabbit anti-VSG121 1:50000 polyclonal antibody, rabbit anti-VSGVO2 polyclonal antibody 1:50 and rat anti-HA mAb 1:500 (Roche) were used as primary antibodies. Alexa Fluor 488, Alexa Fluor 568 goat anti-mouse or anti-rabbit 1:1000 (Life Technologies) and Alexa Fluor 647 anti-mouse (Invitrogen) were used as secondary antibodies. Nuclear and kinetoplast DNA was visualized by DAPI (4,6-diamidino-2-phenylindole) (Life Technologies) staining.

Immunofluorescence microscopy for *Tb*SENP expressing cell line was carried out according to standard protocols (13). For super-resolution microscopy, the cells were attached to poly-L-lysine-treated high-precision coverslips (thickness 1 1/2 mm), stained and then mounted onto glass slides. Cells were stained with 1 µg ml−1 DAPI for 10 min and then mounted in Vectashield without DAPI (super-resolution). Primary antisera were mouse anti-myc (NEB, clone 9B11, 1:2,000) and rabbit anti-Pol-I (largest subunit; 1:100 PMID: 27226299). The secondary antibodies were Alexa Fluor conjugated goat antibodies: anti-mouse and anti-rabbit, Alexa Fluor 488, Alexa Fluor 555 Plus or Alexa Fluor 568 (1:1,000). Cells were analysed using a Zeiss LSM980 Airyscan 2. Representative images obtained by super-resolution microscopy correspond to maximum 3D projections by the brightest intensity of stacks of approximately 30 slices of 0.1 μm. Images acquired with the Zeiss LSM980 Airyscan 2 were deconvolved using Airyscan Joint Deconvolution (XYresolution ∼90 nm). More than 100 G1 cells were used per condition for image quantification in two independent biological replicates.

Images from the VSG switching experiments following 6xHA-LacI-*Tb*UBC9 expression were acquired using either a Zeiss LSM 900 confocal microscope equipped with a Plan-Apochromat 63×/1.4 Oil objective and 405, 488, and 561 nm laser lines with the corresponding emission filters, or a Carl Zeiss Axio Observer 7 microscope equipped with a C-Apochromat 63×/1.20 Water objective and the appropriate fluorescence filter sets.

### Switching experiment

TriFlag-*Tb*SENP trypanosomes were diluted to 10^3^ cells/ml and *Tb*SENP expression was induced for 24 h with 1µg/ml doxycycline. After this period of time the inducer was washed out by centrifugation and parasites were resuspended in fresh media without doxycycline. Trypanosomes were allowed to grow for 72 h and analysed by double immunofluorescence assay using mouse anti-VSG221 mAb and rabbit anti-VSG 121 antiserum.

For immunofluorescence-based quantification of VSG switching in the 6xHA-LacI-*Tb*UBC9 cell line, approximately 10^7^ parasites were harvested after 24, 48, or 72 h of induction with doxycycline (1 µg/ml) and processed for immunofluorescence as described above. Surface expression of VSG221 and VSGVO2 was detected using a mouse monoclonal anti-VSG221 antibody (1:250) and a rabbit polyclonal anti-VSG VO2 antibody (1:50), respectively. Parasites were counted using a Nikon 80i fluorescence microscope equipped with a 100× oil-immersion objective (NA 1.40).

### VSG sequencing

Parasites of RNAi-VSG221, RNAi-VSG221-*Tb*SENP and RNAi-VSG221-*Tb*SENPCXA cell lines were diluted to ∼300 cells/ml and plated in 96 well plates (∼30 cells/ well) in presence of 1 µg/ml doxycycline. After 7-8 days, clones resistant to doxycycline were isolated and analysed by immunofluorescence and Western blot. Total RNA was extracted from switchers using TRIzol reagent (Life Technologies) and treated with RQ1 RNase-Free DNase (Promega). Samples were cleaned by chloroform extraction followed by ethanol precipitation. First strand cDNA synthesis was performed with SuperScript II Reverse Transcriptase (Life Technologies) using random hexamers (Macrogen). Total cDNA was used as PCR template to amplify the VSG variants expressed by each clone using a forward primer that matches the splice leader sequence (GACTAGTTTCTGTACTAT) and a reverse primer that matches a conserved region in all VSG variants (CCGGGTACCGTGTTAAAATATATC) as described in http://tryps.rockefeller.edu/. PCR products were purified and sequenced (Macrogen).

### Switching rate determination

About 20 independent cultures of 1ml final volume were inoculated with either 5 or 50 cells of RNAi-VSG221 or RNAi-VSG221-*Tb*SENP cell lines. Cultures were allowed to grow for ∼8 generations and *Tb*SENP overexpression and/or VSG221 RNAi was induced with 1 µg/mL doxycycline. Each culture was immediately plated by distributing 100 µL aliquots in 10 wells of a 96 well-plate and parasites were allowed to grow for 6-8 days before scoring wells with VSG 221 RNAi resistant parasites (29). Because the number of mutations that occur during the growth of parallel cultures has a Poisson distribution, the mean number of mutations can be calculated using the P_0_ method. In this experiment P_0_-the zero term of this Poisson distribution-is the proportion of cultures with no switchers, which is calculated by counting the number of cultures that do not contain wells with parasites resistant to VSG 221 RNAi, divided by the total number of cultures. The P_0_ value was then used to calculate m (the number of mutations per culture) as m=-ln P_0_. The switching rate was calculated from m/Nt, being Nt the number of parasites in the cultures at the moment of doxycycline addition.

### HaloTag Labeling

Parasites expressing the RNA Pol I–HaloTag fusion were labeled by incubating 10^7^ parasites with either JFX554 HaloTag ligand (150 nM) or JF552 HaloTag ligand (500 nM) for 30 min at 37 °C. Samples were subsequently fixed with 7% formaldehyde in PBS. Fixed parasites were adhered to pretreated microscope slides (Superfrost PlusTM Microscope Slides, VWR) overnight at 4 °C or for 1 h at room temperature, followed by washing with PBS. Immunofluorescence was subsequently performed as previously described in the corresponding section above. Image acquisition was performed on a Carl Zeiss LSM 980 confocal microscope equipped with an Airyscan 2 detector and a C Plan-Apochromat 63×/1.4 Oil DIC M27 objective. Fluorophores were excited using diode laser lines at 405, 488, 561, and 640 nm. Super-resolution imaging was performed using the Airyscan 2 detector in SR mode. Emission filters were selected as appropriate for each fluorophore, and z-stack acquisition was conducted using a pixel size of 0.04 μm and a z-step of 0.15 μm. Airyscan image processing was carried out using ZEN Blue (v3.6), and image deconvolution was performed using the Airyscan Joint Deconvolution algorithm in Dense mode with default settings. Chromatic aberration was corrected using the channel alignment function and FocalCheck fluorescence microscope test slide #1, 1μm beads (Life Technologies) were used to determine channel alignment. Processed images were subsequently analyzed using Fiji.

### Flow Cytometry

Approximately 2.5×10^6^ parasites were harvested by centrifugation at 1000 g for 10 min at room temperature. Parasites were fixed in TDB supplemented with 20 mM glucose and 1% paraformaldehyde (PFA) for 30 min. Following a wash with PBS, cells were incubated with mouse anti-VSG221 mAb (1:100) in PBS containing 1% BSA for 1 h at room temperature under agitation. After washing with PBS, parasites were incubated with Alexa Fluor 488 anti-mouse (1:1000) (Life Technologies) for 1 h at room temperature under agitation, followed by a final wash with PBS. Samples were analyzed using a BD Biosciences LSRFortessa X-20 flow cytometer, and data was processed using FlowJo software (FlowJo LLC, Ashland, OR, USA).

### RNA Extraction and RT–qPCR

RNA extraction and cDNA synthesis was performed as previously described. Real-time quantitative PCR (qPCR) was subsequently performed on an CFX Opus 96 Real Time PCR System (Biorad) with SensiFAST SYBR Lo-ROX with primers for VSG221 (Fw: AGCTAGACGACCAACCGAAGG; Rv: CGCTGGTGCCGCTCTCCTTTG), VSGVO2 (Fw: CAGCGGCTGTACATACTAACA; Rv: TTCAGTTGCTGTGCTTTCGT), *Tb*SENP (Fw: GTCACTCAAGAGCGGAATAG; Rv: GCCAGCCCTCACATTATAC), 7SL (Fw:TGACTTGGTGTTCTGCTTGG; Rv: GTCCGTTGACGGAATCAACC) and C1 (Fw: TTGTGACGACGAGAGCAAAC; Rv: GAAGTGGTTGAACGCCAAAT). Samples and their corresponding dilutions were analyzed in triplicate. Samples processed in the absence of reverse transcriptase were used as controls for residual genomic DNA contamination, whereas no-template controls (water controls) were included to monitor reagent contamination. The results were analyzed using LinReg.

## Acknowledgments

We thank Calvin Tiengwe for providing the HNI cell line, James Budzak for LacO constructs and Gloria Rudenko for the V02 antibody. This work was supported by the National Agency for the Promotion of Science and Technology, Argentina (PICT Grant to VEA 2019-02900) and the Secretariat for Research, Development and Innovation (Grant to VEA 80020250100107SM). JRCF was was supported by a Wellcome Trust/Royal Society Sir Henry Dale Fellowship (222573/Z/21/Z). WH was supported by a PhD scholarship by the Medical Research Council (MRC) DiMeN doctoral programme (MR/W006944/1). The authors used ChatGPT (OpenAI) solely to assist with language editing and improve the clarity of the manuscript. All scientific content, interpretation, and conclusions were developed, reviewed, and approved by the authors.

## Author Contributions

MAB, PAI, MN, JRCF, VEA designed research; MAB, PAI, MS, JRCF performed research; VEA, MAB, MS, MN, JRCF, WH contributed new reagents/analytic tools; MAB, MS, PAI, JRCF, VEA analyzed data; and MAB, MS, PAI, JRCF, VEA wrote the paper.

## Classification

Biological Sciences, Microbiology

## Supplementary material

**Supplementary Figure 1.**
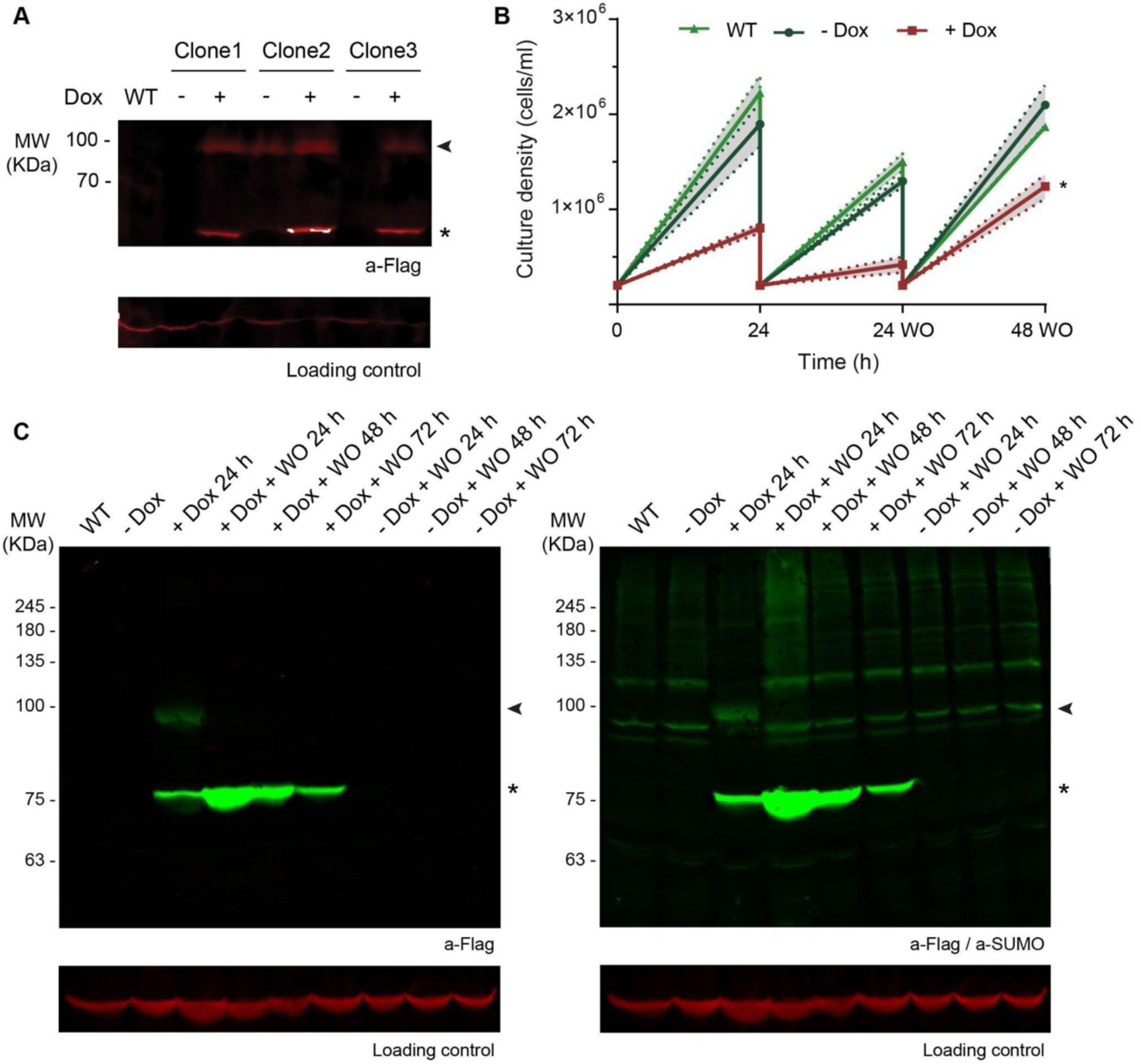
Characterization and validation of doxycycline-inducible ectopic *Tb*SENP-expressing BSF cell lines. **(A)** *Tb*SENP expression in BSF parasites after 24 h induction with doxycycline. Whole-cell extracts of three different clones were analysed by Western blot using anti-Flag antibodies. The arrowhead indicates the full-length protein, while the asterisk indicates a lower-molecular-weight form. Tubuline was used as loading control. **(B)** Growth curves of wild-type (WT), uninduced (-Dox), and induced (+Dox) *Tb*SENP parasites, alongside subsequent 24 h and 48 h washout (WO) time points. Statistical analysis: Student’s t-test, *P<0.05. **(C)** Analysis of *Tb*SENP expression after 24 h of induction with doxycycline, alongside subsequent 24 h, 48 h and 72 h washout (WO) time points. Uninduced parasites (-Dox) were used as a control. Whole-cell extracts were analyzed by Western blot using anti-Flag antibodies for *Tb*SENP detection (left panel) and anti-*Tb*SUMO antibodies for *Tb*SUMO detection (right panel). The arrowhead indicates the full-length protein, while the asterisk indicates a lower-molecular-weight form. PABP-C was used as a loading control.

**Supplementary Figure 2.**
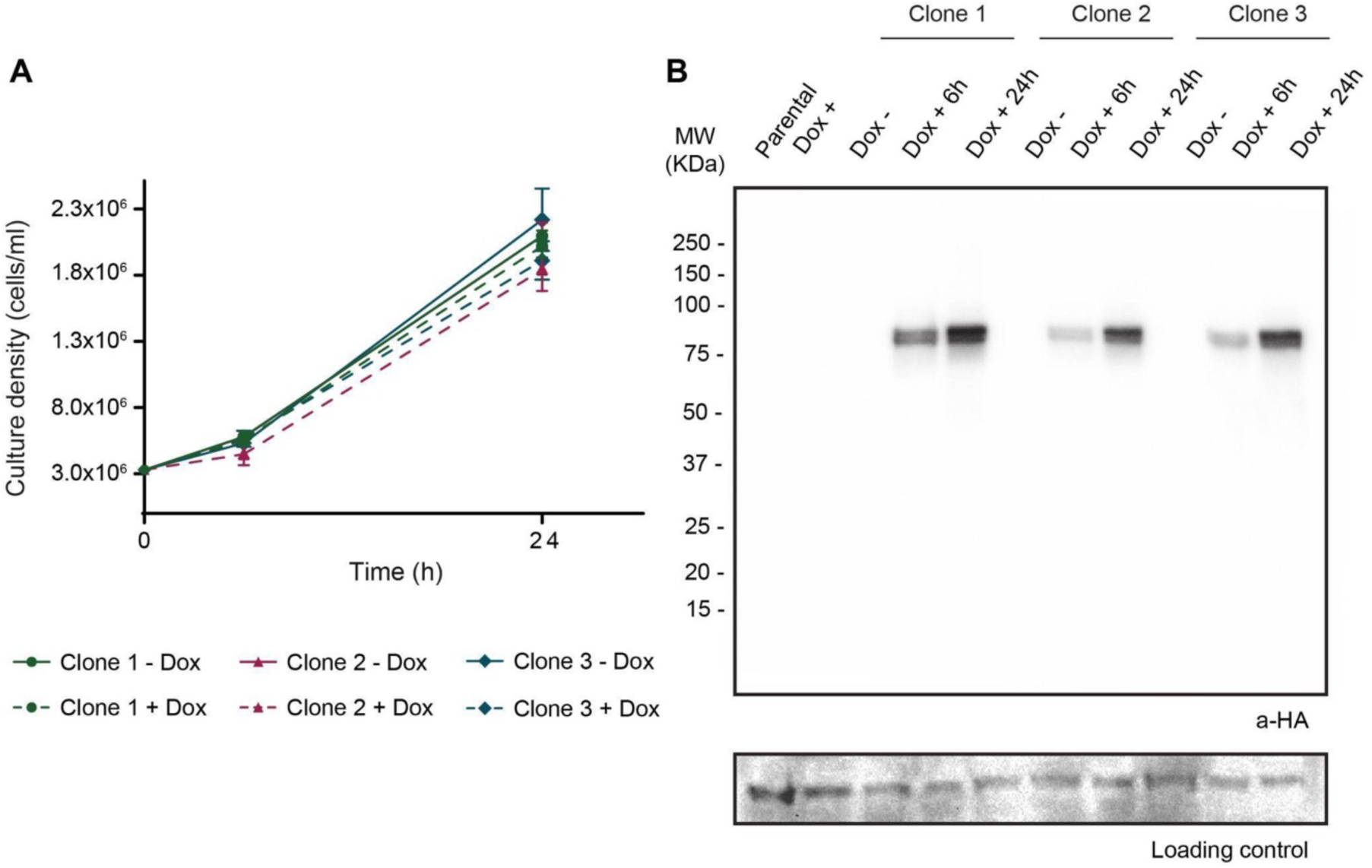
Validation of doxycycline-inducible 6XHA-LacI-*Tb*UBC9 and BES1-repeats expressing BSF cell lines. **(A)** Growth curves of uninduced (-Dox), and induced (+Dox) 6XHA-LacI-*Tb*UBC9 clones. Error bars show standard deviation. Statistical analysis: multiple Student’s t-test, *P<0.05. **(B)** 6XHA-LacI-*Tb*UBC9 expression in BSF parasites after 6 h and 24 h induction with doxycycline. Whole-cell extracts of three different clones were analysed by Western blot using anti-HA antibodies. BIP was used as loading control.

**Supplementary Figure 3.**
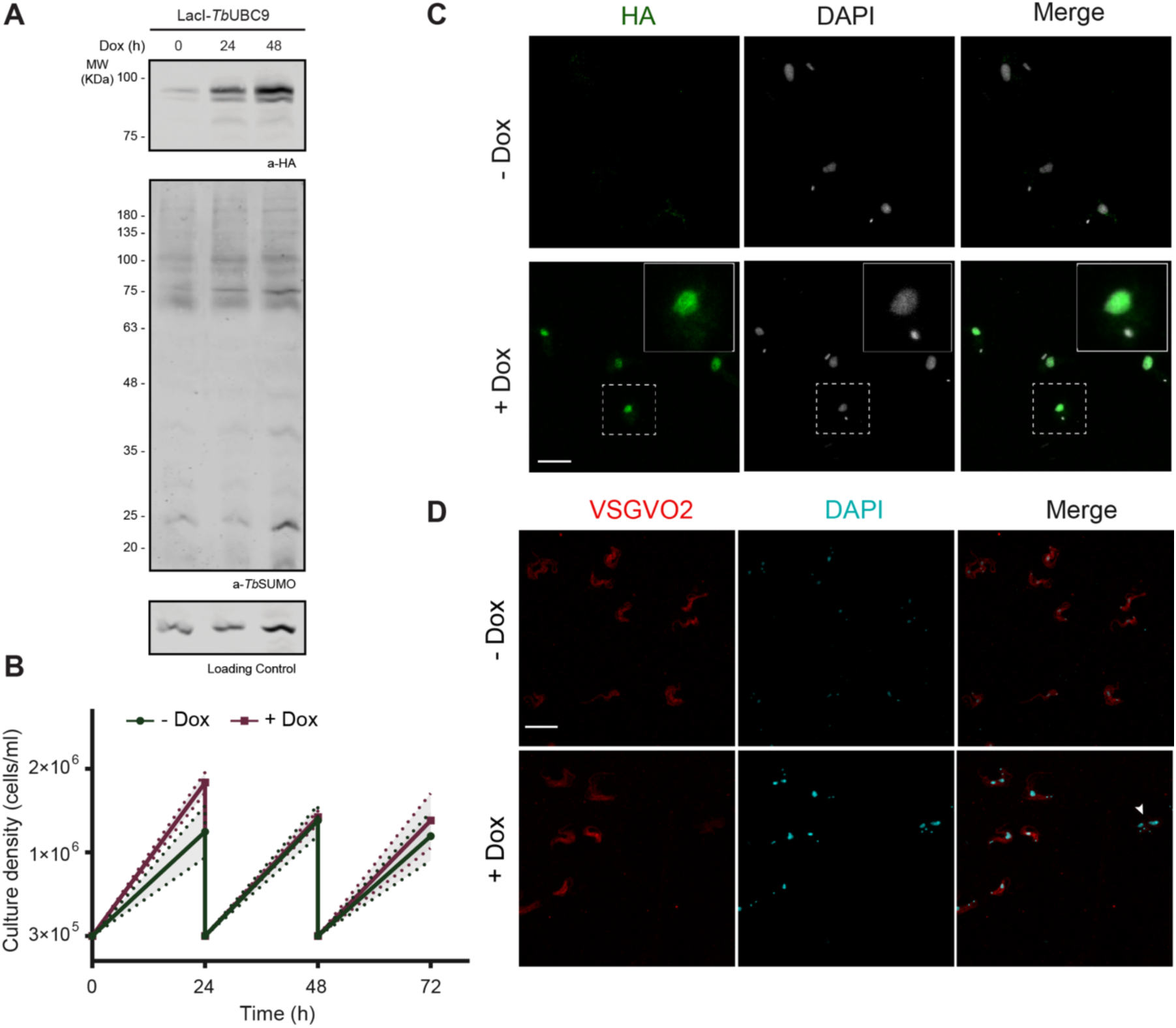
Characterisation of doxycycline-inducible 6XHA-LacI-*Tb*UBC9 expressing BSF cell lines in the absence of BES1-LacO-repeats. **(A)** Western blot analysis of 6XHA-LacI-*Tb*UBC9 parasites lacking lacO repeats under non-induced (Dox 0 h) and induced conditions (Dox 24 h, 48 h). Whole cell lysates were probed with anti-HA and anti-*Tb*SUMO antibodies. Actine was used as loading control. **(B)** Growth curves of 6XHA-LacI-*Tb*UBC9 parasites lacking lacO with (+Dox) or without (-Dox) doxycycline maintained with 24 hourly subculture (n=3 independent induction). Statistical analysis, multiple Error bars show standard deviation. Statistical analysis: multiple Student’s t-test *P<0.05. **(C)** Indirect immunofluorescence of 6XHA-LacI-*Tb*UBC9 parasites lacking lacO repeats using anti-VSGVO2 antibodies (red). Representative images of the population with (+Dox) and without (-Dox) doxycycline induction are shown. Arrowhead indicates VSGVO2 negative parasites. Nuclei and kinetoplasts were stained with DAPI (cyan). Scale bar: 10 μm.

**Supplementary Figure 4.**
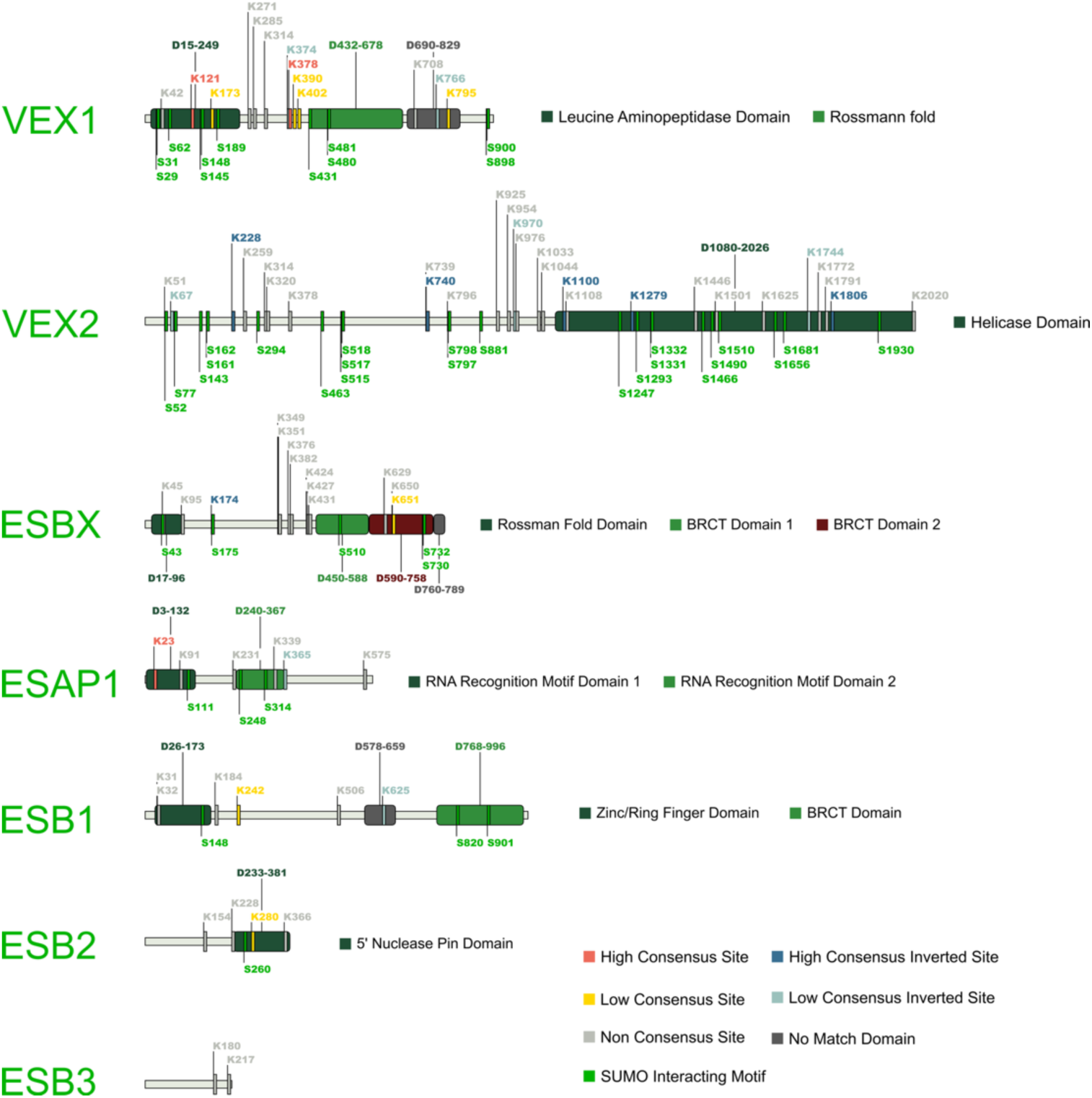
Various *T. brucei* ESB residing or associated proteins show multiple predicted SUMOylation sites and SUMO interacting motifs. Protein maps were generated through a custom-made Python script, protein domains were identified through a combination of AlphaFold3 (54) and Foldseek (55) analyses, and JASSA (56) was used for predicting SUMOylation and SUMO interacting motifs (SIM) sites. Protein maps are scaled to total amino acid length. Regarding predicted SUMOylation sites: consensus sites are short sequences of amino acids which have been shown in at least one study to be SUMOylated; high consensus sites represent regularly SUMOylated areas, whilst low consensus sites are less commonly SUMOylated; non consensus sites have been shown to be SUMOylated at least once; lastly, the inverted consensus sites’ amino acid sequences are flipped in relation to normal consensus sites. SIMs describe sites where SUMO interacts with proteins non-covalently.

---

**Supplementary Table 1.** Calculation of switching rates following *Tb*SENP overexpression. The initial parasite density (N0) and the parasite density at the time of VSG221 RNAi induction (Nt) are indicated as parasites/ml. The number of cultures analyzed is indicated as C. Cultures were distributed into 96-well multiwell plates following doxycycline induction. The fraction of cultures lacking parasites resistant to VSG221 RNAi is indicated as P_0_. The mean number of VSG RNAi-resistant cells per culture is indicated as M. The antigenic variation rate per cell generation (µ) is shown in the last row.

|  | **RNAi VSG221**<br>5 cells/ml | **RNAi VSG221**<br>50 cells/ml | **RNAi VSG221-*Tb*SENP**<br>5 cells/ml | **RNAi VSG221-*Tb*SENP**<br>50 cells/ml |
| --- | --- | --- | --- | --- |
| **N<sub>0</sub>** | 5 | 50 | 5 | 50 |
| **N<sub>t</sub>** | 3750 | 33500 | 1612 | 15000 |
| **C** | 8 | 10 | 9 | 10 |
| **P<sub>0</sub>** | 0.62 | 0.20 | 0.11 | - |
| **M** | 0.48 | 1.61 | 2.20 | - |
| **Rate** | $1.28 \times 10^{-4}$ | $4.80 \times 10^{-5}$ | $1.36 \times 10^{-3}$ | - |

